# From Resonance to Computation: A Six-Layer Framework for Analog Neural Processing in Coupled RLC Oscillator Networks

**DOI:** 10.64898/2026.04.09.717435

**Authors:** Jeremy Sender

**Affiliations:** Department of Computer Science and Information Technology, La Trobe University, Melbourne, Australia

**Keywords:** Analog computation, neural resonance, RLC circuit, attractor dynamics, phase coding, subthreshold impedance, neuromorphic engineering, coupled oscillators

## Abstract

Subthreshold neuronal membranes exhibit resonant, band-pass impedance characterised by an effective inductance arising from voltage-gated channel kinetics—principally *I_h_*. This paper presents a six-layer computational framework that builds from this single-neuron RLC description to a complete account of how coupled neural oscillator networks compute. Layer 1 establishes the RLC neuron as a frequency-selective bandpass filter. Layer 2 shows that coupled RLC neurons encode relational information in phase differences (binding). Layer 3 demonstrates that networks of coupled oscillators form attractor landscapes supporting memory and pattern completion, with fixed-point, limit-cycle, and chaotic attractor classes. Layer 4 identifies the synaptic coupling matrix as a learned impedance network whose topology determines attractor geometry. Layer 5 maps neuromodulatory systems to bias controls that sweep RLC parameters (resonant frequency, quality factor, gain) without modifying stored memories. Layer 6 assembles the full system with cross-frequency multiplexing and homeostatic stabilisation. The framework is grounded in measurable electrical quantities and generates testable predictions distinguishing it from rate-coding and RC integrate-and-fire models. We explicitly address the linearisation gap between the subthreshold regime where the RLC description is rigorous and the nonlinear regime where attractor dynamics operate, the noise and precision limits of analog neural computation (∼ 3.3 effective bits per neuron, compensated by massive parallelism), and the distinction between causal and correlative evidence for oscillation-based coding claims. The framework does not replace existing models; it extends them by showing that rate coding is one (coarse) description of the output of an analog computation whose richer dynamics— resonance, phase, temporal fine structure—may carry additional computational content.

## I. Introduction

Neurons are electrical circuits. This observation dates to Hodgkin and Huxley [10], who drew the neuronal membrane as a circuit schematic—capacitor in parallel with voltage-dependent conductances and ionic batteries—and it remains the foundation of computational neuroscience seven decades later. What has received less systematic attention is the consequence: if neurons are circuits, then the tools of analog electrical engineering—impedance spectroscopy, transfer functions, Bode plots, stability analysis, noise theory—should be directly applicable to understanding neural computation.

A growing body of evidence supports this perspective. Subthreshold resonance, mediated by the hyperpolarisation-activated cation current *I*_*h*_, has been documented in hippocampal pyramidal cells, entorhinal stellate cells, thalamic relay neurons, inferior olivary neurons, and cortical interneurons [15]. The linearised subthreshold impedance of these neurons is well-described by a parallel RLC circuit, with effective inductance arising from the delayed kinetics of *I*_*h*_ [21]. This is not metaphor: the impedance can be measured with sinusoidal current injection, the resonant frequency *f*_0_ read from the impedance peak, and the quality factor *Q* estimated from the peak width.

In a companion paper [33], we catalogued fifteen neural circuits whose topology and transfer function match specific, named analog electronic designs—from the basilar membrane as a distributed *LC* filter bank to central pattern generators as astable multivibrators to gap junctions as resistive coupling. That catalogue established the component-level correspondences. The present paper addresses the question those correspondences leave open: *how do these circuit primitives compute when coupled into networks?*

The answer is developed through six layers, each building on the previous:

1. **Layer 1:** The single RLC neuron as frequency-selective bandpass filter.
2. **Layer 2:** Coupled RLC neurons encoding relational information in phase differences.
3. **Layer 3:** Small networks forming attractor landscapes for memory and categorisation.
4. **Layer 4:** The synaptic coupling matrix as a learned impedance network.
5. **Layer 5:** Neuromodulation as parameter sweep (bias control) of RLC circuit values.
6. **Layer 6:** The full system with cross-frequency multiplexing and homeostatic stabilisation.

The lower layers rest on experimentally validated biophysics; the upper layers are increasingly theoretical. Sections VIII–X address where the framework is rigorous, where it becomes heuristic, and where evidence is causal versus correlative.

The framework does not replace rate-coding or integrate-and-fire models. It extends them: rate is one description of the output of an analog computation whose richer dynamics— resonance, phase, temporal fine structure—carry additional computational content that rate-based models discard.

## II. Layer 1: The Single RLC Neuron as Signal Processor

A single neuron at subthreshold potentials, linearised around rest, has an impedance taking the form of a parallel RLC resonance [15], [21]:

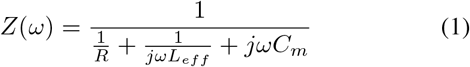

with resonant frequency 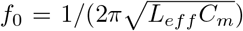 and quality factor 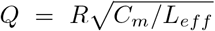, where *R* = 1*/g*_*L*_ is the leak resistance, *C*_*m*_ is the membrane capacitance, and *L*_*eff*_ is the effective inductance arising from the delayed kinetics of voltage-gated channels—principally the hyperpolarisation-activated cation current *I*_*h*_.

The effective inductance is not a physical inductor. It is a phenomenological quantity that captures the phase-lead behaviour produced by the voltage-dependent kinetics of *I*_*h*_: when the membrane depolarises, *I*_*h*_ deactivates with a time delay, producing a current that opposes the initial voltage change with a phase relationship equivalent to an inductive element [25]. The explicit expression, derived from linearisation of the Hodgkin–Huxley equations, is:

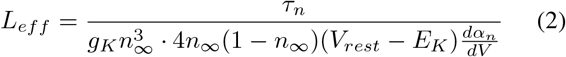

where the variables follow standard Hodgkin–Huxley notation.

### The critical distinction from integrate-and-fire models

An RC neuron (*L*_*eff*_ → ∞, no inductive term) is a lowpass filter—it integrates all inputs equally, attenuating high frequencies. An RLC neuron is a *bandpass* filter—it selectively amplifies inputs near *f*_0_ and attenuates inputs far from *f*_0_. The gain at resonance relative to the DC response is:

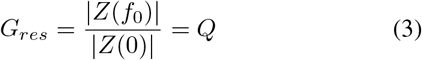

For a neuron with *Q* = 5 (hippocampal pyramidal cells in the theta band), inputs at *f*_0_ are amplified 5× relative to DC. For *Q* = 10× (entorhinal stellate cells), the amplification is 10×. This selective amplification is invisible to RC models.

### Experimental evidence

Subthreshold resonance has been measured across cell types: hippocampal pyramidal cells (*f*_0_ ≈ 4–8 Hz), entorhinal stellate cells (*f*_0_ ≈ 8–12 Hz), inferior olivary neurons (*f*_0_ ≈ 3–10 Hz), thalamic relay neurons (*f*_0_ ≈ 2– 10 Hz), and cortical interneurons (*f*_0_ ≈ 20–50 Hz) [15]. In each case, resonance is abolished by *I*_*h*_ blockers (ZD7288, Cs^+^), confirming the causal role of *I*_*h*_ [15]. The framework’s equations are agnostic to the molecular identity of the current producing *L*_*eff*_ —any delayed voltage-dependent conductance with appropriate kinetics suffices. In thalamocortical neurons, the low-threshold calcium current *I*_*T*_ rather than *I*_*h*_ generates resonance [35]; the RLC description applies equally, with *L*_*eff*_ arising from *I*_*T*_ kinetics instead. Independent support comes from tACS studies: Asamoah et al. [2] showed that weak extracellular fields modulate oscillation amplitude following a resonance pattern—large effects at endogenous frequency, negligible effects off-resonance—as the bandpass model predicts and RC models do not.

The RLC description is valid at subthreshold potentials within ∼ 10–15 mV of rest, for small-signal perturbations. At more depolarised or hyperpolarised potentials, *L*_*eff*_ depends on voltage through the gating variables, and at sufficiently hyperpolarised levels the inductive term can vanish, with the impedance degenerating to RC [21]. The framework is therefore state-dependent and is most informative in the near-rest operating regime where neurons spend the majority of their time in vivo.

### Testable prediction

*Neurons with measurable resonance (Q >* 2*) should show frequency-dependent spike-timing precision: stimuli at f*_0_ *should produce more precisely timed spikes (lower jitter) than stimuli at non-resonant frequencies. This has been confirmed in entorhinal stellate cells [9]; analogous experiments in inferior olivary and thalamic relay neurons would provide independent tests*.

## III. Layer 2: Coupled RLC Neurons—Phase Relationships

Connect two RLC neurons with coupling conductance *g* (resistive for gap junctions, frequency-dependent for chemical synapses). The coupled system is:

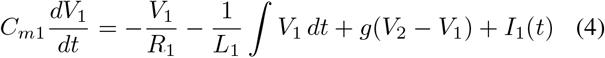

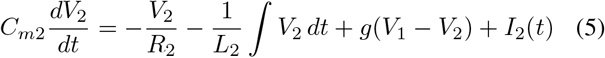

Two regimes emerge depending on coupling strength *g* relative to the frequency mismatch Δ*f* = |*f*_01_ − *f*_02_|:

### Strong coupling (*g* ≫ 2*π*Δ*f* · *C*_*m*_)

The oscillators phase-lock to a common frequency with a fixed phase difference:

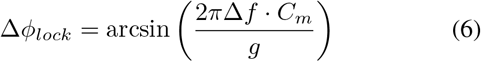

This phase difference is a continuous, measurable quantity encoding the frequency mismatch between the two neurons. Phase locking is a candidate mechanism for *binding*—two neurons with locked phase are “bound” into a functional assembly. The evidence for this interpretation is convergent across multiple systems but predominantly correlative rather than fully causal (see the evidence grading in Section X).

### Worked example

Consider two hippocampal CA1 pyramidal neurons: *f*_01_ = 6 Hz, *f*_02_ = 7 Hz, *C*_*m*_ = 200 pF, coupled by a gap junction with *g*_*gap*_ = 1 nS. The frequency mismatch is Δ*f* = 1 Hz. The coupling threshold for locking requires *g >* 2*π* · 1 · 200 × 10^−12^ ≈ 1.26 nS. With *g*_*gap*_ = 1 nS, we are below threshold: the neurons show intermittent phase slipping. With a second gap junction (total *g* = 2 nS *>* 1.26 nS), they lock with Δ*ϕ* = arcsin(1.26*/*2.0) = arcsin(0.63) ≈ 39. This 39° offset continuously encodes the 1 Hz mismatch. Increasing to *g* = 5 nS gives Δ*ϕ* ≈ 14.5—stronger coupling pulls phases together, as expected from Equation 6.

**Fig. 1.**
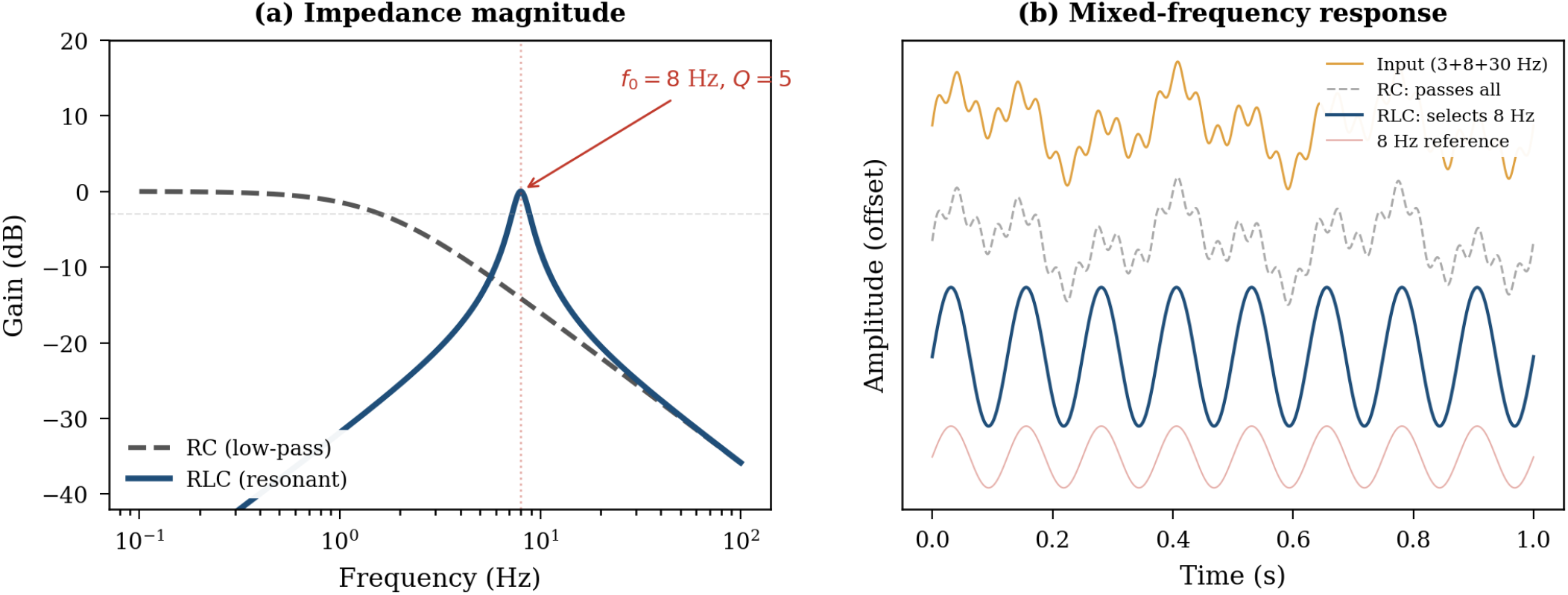
RC vs RLC neuron frequency response. Left: Bode plot showing the RC neuron’s flat low-pass characteristic versus the RLC neuron’s resonant peak at ∼ 8 Hz (theta range) with *Q* = 5. Right: Response to mixed-frequency input (3 + 8 + 30 Hz). The RC neuron passes all components; the RLC neuron selectively extracts the 8 Hz component.

### Weak coupling (*g* ≪ 2*π*Δ*f* · *C*_*m*_)

The oscillators do not lock. Their superposition produces amplitude modulation at the beat frequency:

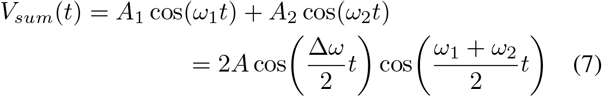

When *f*_1_ ≈ 40 Hz (gamma) and *f*_2_ ≈ 6 Hz (theta), the beat produces theta-modulated gamma bursts, consistent with hippocampal cross-frequency coupling [17].

### What is computed

Relational encoding. Zero phase difference = correlated inputs (bound); *π* phase = anti-correlated (competing); random phase = uncorrelated (independent). This information resides in the *relationship* between neurons, not in either neuron’s rate. Phase codes require a stable reference— typically provided by population oscillations periodically reset by salient events—and the framework assumes such references are available (see the causal/correlative grading in Section X). Operationally, a phase reference can be detected via event-locked phase reset: the instantaneous phase of the LFP theta oscillation is computed (e.g., via Hilbert transform), and reset events are identified as time points where phase variance across trials drops below a threshold. In hippocampal recordings, such resets occur reliably at place-field entry, sharp-wave ripple onset, and sensory-evoked theta reset. In the model, a reference oscillator population with strong internal coupling and weak external drive provides a stable clock against which individual neuron phases are measured; the coupling equations of Layer 2 then describe phase offsets relative to this reference.

## IV. Layer 3: Small Networks—Attractor Formation

We now move from pairwise phase relations to collective computation in small recurrent networks, where memory and categorisation arise from the geometry of state space and from convergence—or structured wandering—under recurrent coupling.

### A. Network Dynamics

Consider *N* resonant elements whose subthreshold state is represented by a voltage-like variable *v*_*i*_(*t*) and a conjugate reactive variable capturing second-order dynamics, coupled through a weighted adjacency matrix *W* = [*w*_*ij*_]. A minimal second-order network equation retaining the key RLC feature (reactive inertia plus dissipation) is:

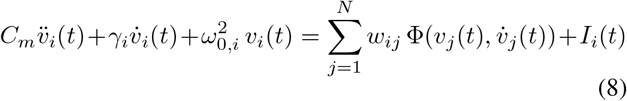

where *C*_*m*_ is the effective capacitance, *γ*_*i*_ is a damping term, *ω*_0,*i*_ = 2*πf*_0,*i*_ sets the intrinsic resonant timescale, *I*_*i*_(*t*) is external drive, and Φ( · ) is a coupling function that may be approximately linear (subthreshold) or nonlinear and state-dependent (spike-mediated). This form accommodates retrieval via convergence to a static pattern, a dynamical orbit, or a chaotic regime.

### B. Energy-Like Functions and the Lyapunov Gap

For first-order attractor models, an energy function guarantees convergence and yields sharp capacity predictions [11]. For second-order dynamics, we motivate an energy-like decomposition into kinetic, potential, and coupling terms:

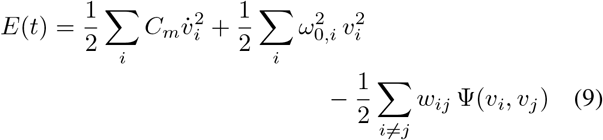

The three terms represent kinetic, potential, and coupling contributions respectively. Ψ is chosen so that phase-consistent or pattern-consistent configurations lower the coupling term. Oscillatory transients correspond naturally to exchange between kinetic and potential contributions rather than to monotone descent at every instant.

**Fig. 2.**
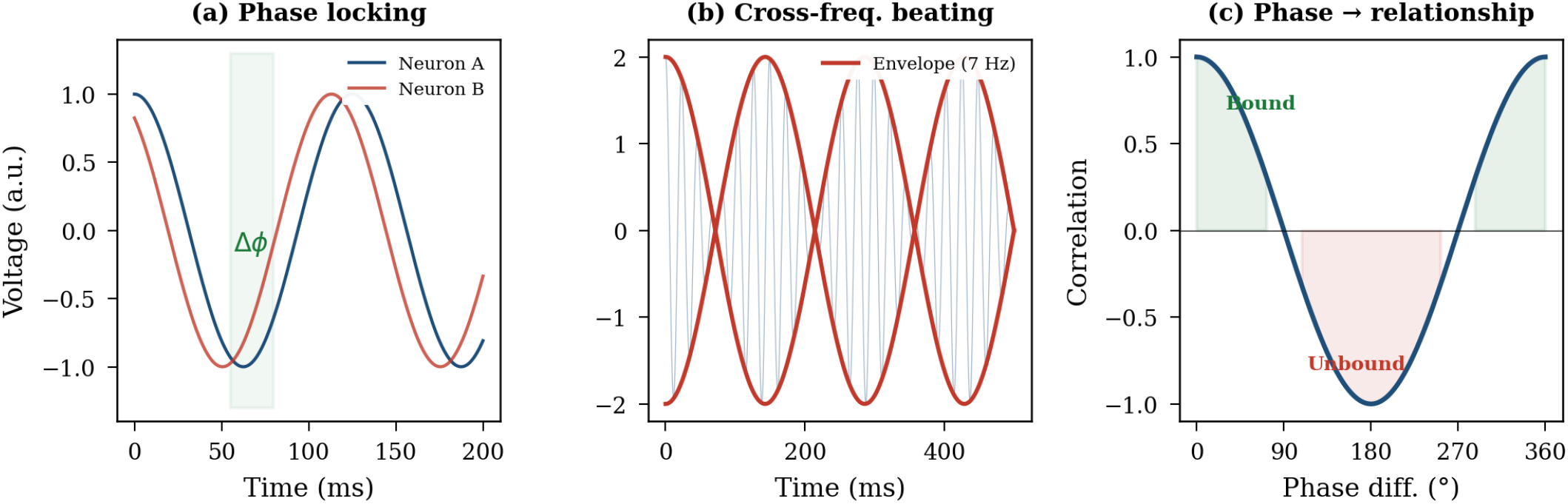
Coupled oscillator dynamics. Left: Phase locking under strong coupling. Centre: Cross-frequency beating under weak coupling. Right: Phase difference encodes input correlation.

The theoretical gap is straightforward to state: although *E*(*t*) can be computed and relaxation-like behaviour is observed in damped regimes, local stability can coexist with global instability, and energy-like intuition should not be taken to imply global convergence without proof.

#### Proposition 1

*(Sufficient conditions for energy descent):* Consider the network (8) with:

i. symmetric coupling: *w*_*ij*_ = *w*_*ji*_ for all *i, j*;
ii. linear coupling function: 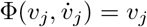
iii. uniform positive damping: *γ*_*i*_ ≥ *γ*_*crit*_ *>* 0 for all *i*,

where *γ*_*crit*_ depends on the spectral radius *ρ*(*W* ) and the spread of natural frequencies {*ω*_0,*i*_ }. Under these conditions, *dE/dt* ≤ 0 along all trajectories, the network reduces to a dissipative gradient system, and convergence to local minima of *E* is guaranteed.

For the underdamped regime (*γ*_*i*_ *< γ*_*crit*_, equivalently *Q > Q*_*crit*_), transient oscillations occur and monotone descent fails at individual time steps, but numerical simulations of networks up to *N* = 100 show convergence after transient ringing. Whether *E* is a strict Lyapunov function for the general underdamped, nonlinear-Φ case remains open. Prior work provides a benchmark: Aoyagi [1] proved Lyapunov stability for phase-coded oscillator memories under specific coupling symmetry; the present framework’s claim is that RLC-grounded parameters constrain the same class of dynamics, not that we have extended the proof.

In practice, the biologically relevant criterion is not monotone descent but whether the network settles into a reliable attractor regime on a physiological timescale despite transient ringing and noise. This motivates using tolerance-based settling times *τ* (*η*) as the evaluation metric rather than asymptotic convergence proofs, since convergence times can exhibit heavy-tailed behaviour depending on load and initial overlap—retrieval latency is itself a meaningful computational variable.

### C. Capacity, Transient Retrieval, and Operational Success

Classical analyses establish a sharp capacity threshold: error-free retrieval up to a storage limit, catastrophic failure beyond it [7], [11]. In oscillatory networks with STDP-based phase-coded learning, Scarpetta et al. [30] found capacity scaling linearly with *N*, suggesting that phase degrees of freedom may expand storage beyond the classical bound. Modern dynamical analyses further show that useful computation can occur above the classical threshold when one adopts a finite-time notion of “usable retrieval”: patterns may be transiently retrieved with high accuracy even in the absence of stable attractors.

We define success using a hierarchy: *recognition* as the satisfaction of necessary stability conditions for a target pattern; *recall* as convergence (or entry into a bounded dynamical regime) within a tolerance tube around the target for at least *τ* (*η*); and *transient retrieval* as the ability to approach the target with high accuracy for a task-relevant window even when the state later departs. This framing naturally distinguishes failures due to confusion among learned items from failures due to genuinely spurious attractors.

### D. Beyond Fixed Points: Limit Cycles and Chaos

A strict fixed-point view is inadequate for biological settings where learning is continuous and forgetting is unavoidable. When new memories are learned and old memories decay under an online synaptic rule, retrieval dynamics become age-dependent: recent memories are retrieved as fixed-point attractors while older memories are retrieved as chaotic attractors characterised by heterogeneity and temporal fluctuations. Fixed-point and chaotic attractors coexist in the same phase space, with chaotic fluctuations increasing and basins shrinking as memories age—a natural mechanism by which older memories become less robust even if they remain partially retrievable.

This supports a three-class taxonomy. **Fixed-point attractors** implement classical content-addressable recall and pattern completion [11]. **Limit-cycle attractors** implement rhythmic or phase-structured memories, where retrieval corresponds to convergence onto a periodic orbit [20]; partial presentation of a phase-consistent cue can induce selective retrieval provided storage is within capacity [31]. **Strange (chaotic) attractors** implement bounded but non-repeating retrieval regimes where local stability coexists with global instability [8]. The effective inductance introduces the second-order dynamics that permit oscillatory and chaotic solutions forbidden in first-order (RC) networks; the transition between attractor types can be driven by neuromodulatory parameter changes (Section VI).

**Fig. 3.**
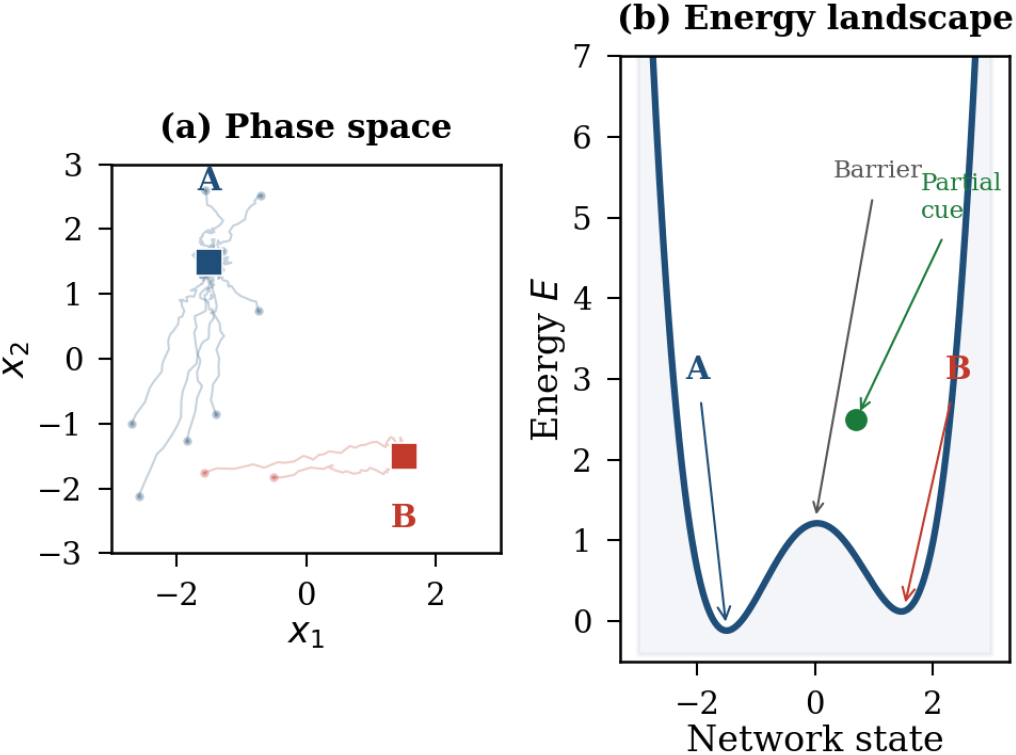
Attractor dynamics. Left: Trajectories converge to one of two point attractors (stored memories). Right: Energy landscape with basins separated by a barrier; a partial cue rolls to the nearest minimum (pattern completion).

### E. Relationship to Prior Work

The importance of phase variables for robust recovery is supported by evidence that phase information can outperform amplitude information in reconstructing incomplete or corrupted stimuli. Aoyagi [1] proved Lyapunov stability for phase-coded oscillator memories, and Scarpetta et al. [30] demonstrated STDP-based storage of phase-coded spatiotemporal patterns with linear capacity scaling. The RLC framework grounds these results in measurable neuron-level quantities: the resonant frequency *f*_0_ and quality factor *Q* that determine achievable phase relationships are set by channel kinetics and membrane capacitance, not free parameters.

## V. Layer 4: The Coupling Matrix as Learned Impedance Network

At Layer 4, we interpret the recurrent coupling matrix *W* not merely as abstract synaptic weights but as a learned impedance structure governing which phase relations are stable, which assemblies can synchronise, and which attractor basins exist in practice. Learning rules that preferentially strengthen coherent phase relations sculpt the set of stable assemblies the network can express.

### A. A Phase-Based Hebbian Rule

We adopt a continuous-time phase-Hebbian update:

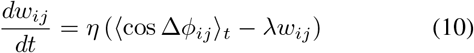

where Δ*ϕ*_*ij*_(*t*) is the instantaneous phase difference between nodes *i* and *j, η* sets a learning rate, and *λ* implements homeostatic weight decay. In-phase pairs are reinforced; anti-phase pairs are weakened, making synchronised assemblies easier to re-enter and more robust to noise. Phase-coupling rules of this form exist in the coupled-oscillator literature [12], [13]; what the RLC framework adds is that the phase relationships being learned are constrained by measurable impedance properties (*f*_0_, *Q*) of the constituent neurons.

This continuous cosine rule should be contrasted with spike-timing-dependent phase-coded learning, where Scarpetta and Giacco [31] showed that capacity scales linearly with *N* and retrieval frequency depends on the STDP time window asymmetry. Whether continuous subthreshold phase-Hebbian updates produce comparable capacity scaling and basin geometry to spike-window STDP updates in the same noise regime is an open question connecting the RLC framework to the phase-coded memory literature.

### B. Landscape Quality and Training Objectives

Naive Hebbian learning can produce spurious attractors and shallow, overlapping basins. The dominant failure mode at low-to-moderate loads is not convergence to a spurious state but confusion: convergence to an incorrect but valid learned pattern. Under extreme high-load and high-noise conditions, spurious errors become significant and landscape pathologies re-emerge. Improving *W* is therefore not only a matter of increasing nominal capacity but of reshaping the error spectrum across operating regimes—confusion among learned patterns, convergence to spurious states, and failure to converge within a physiological time budget. Convergence-time statistics, including power-law scaling with load, should be treated as part of the evaluation criteria rather than as a binary asymptotic property.

### C. Spectral Structure

The topology of *W* determines attractor geometry. The graph Laplacian ℒ = *D* − *W* governs the spectral properties of the network: its eigenvalues determine attractor basin separation and convergence timescales. Synchronisation stability and cluster structure depend on how coupling is distributed across the graph. We highlight as an open mathematical direction whether equilibrium embeddings of spring networks (spring constants ∝ *w*_*ij*_, determined by ℒ) provide a useful analogy for predicting attractor locations from the spectrum of the learned impedance network.

## VI. Layer 5: Neuromodulation as Bias Control

Neuromodulators reshape intrinsic and network-level dynamical parameters—frequency preference, resonance strength, coupling efficacy, effective stability—without being the content-bearing signal itself. They are *bias controls*: they set the operating point. The expression of intrinsic-current contributions depends on whether synaptic transmission is intact, so bias-control manipulations can have different consequences in active versus quiescent networks.

### A. Serotonin and I_h_

Serotonin illustrates why bias control must be formulated as receptor- and context-dependent. Bath application of 5-HT suppresses *I*_*h*_-dependent sag and conductance with modest shifts in activation voltage dependence [22], [26]. Low-frequency resonance below resting potential is reduced, directly implying that serotonergic *I*_*h*_ modulation can alter frequency preference to synaptic inputs. The signalling pathway matters: suppression involves cAMP/PKA rather than PKC, and effect size differs across developmental stages— bias control is a physiological state variable, not a fixed knob.

Receptor subtype provides a mechanistic axis for bidirectionality. Activating 5-HT_1*A*_ receptors reduces *I*_*h*_ and shifts activation hyperpolarising; activating 5-HT_4_ and 5-HT_7_ receptors increases *I*_*h*_ and shifts activation depolarising. Any mapping from “serotonin” to a single RLC parameter must therefore be understood as an aggregate population statement depending on receptor expression and cellular context.

### B. Acetylcholine and Multi-Current Tuning

Cholinergic modulation directly demonstrates multi-parameter bias control: resonance frequency and resonance strength vary as a function of cholinergic modulation [4], [18], mediated through both *I*_*h*_ and *I*_*M*_ . At the circuit-behaviour interface, cholinergic agonism preferentially modulates low-frequency alpha/beta oscillatory organisation and correlates with speeded performance, while leaving high-frequency gamma oscillations unaltered—consistent with neuromodulatory re-weighting of frequency channels rather than uniform gain.

### C. Histamine and Impedance-Derived Resonance Metrics

Histamine provides a direct bridge from neuromodulator action to impedance-derived resonance metrics. Using impedance amplitude profiling (ZAP), histamine increases both resonant frequency *f*_*res*_ and its magnitude (a *Q*-like measure) in a concentration-dependent manner resembling histamine’s action on *I*_*h*_. This link is pharmacologically constrained: ZD-7288 blocks both the histamine-induced action and resonance, supporting the inference that *I*_*h*_ is a necessary pathway for this resonance modulation in that preparation.

### D. Dopamine and Attractor Accessibility

Dopamine connects bias control to attractor computation. Phasic dopamine reveals latent network attractors previously built by dopamine-modulated plasticity, promoting engagement into decision-related attractor dynamics [32]. Dopamine also instantaneously potentiates NMDA efficacy, providing a mechanistic pathway by which it can reshape basin depth and transition rates without rewriting stored patterns.

### E. What Is Computed

Mode selection. Same network, same stored memories, different computational character depending on which bias controls are active. As a heuristic mapping between parameter regimes and computational phenotypes: high *Q* corresponds to focused, frequency-selective processing (a correlate of attention); low *Q* to broad integration (a correlate of drowsy or diffuse states); elevated noise to exploratory dynamics. This mapping describes aggregate parameter-regime behaviour, not a one-to-one psychological identification; any given neuro-modulator can have opposing effects via different receptor subtypes (Section VI-A), and the net computational phenotype depends on the receptor mixture active in a given circuit at a given time. The intended claim is correspondingly aggregate: neuromodulators shift populations of resonant elements through families of impedance profiles, thereby biasing the set of phase relations, attractor basins, and convergence times the network can realise within its physiological noise and time budgets.

### F. Context Dependence

A central constraint on Layer 5 is that intrinsic-current effects can be gated by the synaptic regime. In L5 pyramidal networks, driven networks exhibit theta-band resonance with *I*_*h*_ and *I*_*M*_ contributing to suprathreshold network responses. Blocking KCNQ or HCN channels boosts low-frequency responses, demonstrating measurable intrinsic contributions to network-level frequency response. However, when synaptic transmission is suppressed, blocking HCN channels has minimal effect—the expression of *I*_*h*_-dependent resonance at the network level depends on whether synaptic interactions are engaged. Neuromodulation as bias control is therefore a controlled simplification: the receptor-specific bidirectionality of serotonergic *I*_*h*_ modulation, the multi-current mediation of cholinergic resonance changes, and the regime dependence in intact versus synaptically suppressed networks all indicate that neuromodulatory bias control is fundamentally multidimensional and context-sensitive.

## VII. Layer 6: The Full System

Assembling all layers: (1) Individual RLC neurons perform frequency-selective filtering. (2) Coupled pairs encode phase relationships. (3) Networks form attractor landscapes. (4) The coupling matrix stores learned patterns. (5) Neuromodulation tunes the computational mode. (6) Cross-frequency coupling (frequency-division multiplexing) allows multiple computations in parallel on different carrier frequencies [17], while homeostatic plasticity [34] maintains the system within its operating range.

### The spike as VCO readout

The axon hillock functions as a voltage-controlled oscillator: the continuous subthreshold trajectory controls when each spike fires. Rate coding—counting spikes per second—is a lossy compression of this continuous information. The analog framework does not replace rate coding; it extends it: rate is one (coarse) description of the output of an analog computation.

### What “computation” means here

Not instruction execution. Not symbol manipulation. The continuous evolution of a high-dimensional dynamical system through a structured phase space toward stable configurations encoding meaningful relationships in the input. The computation is the trajectory.

**Fig. 4.**
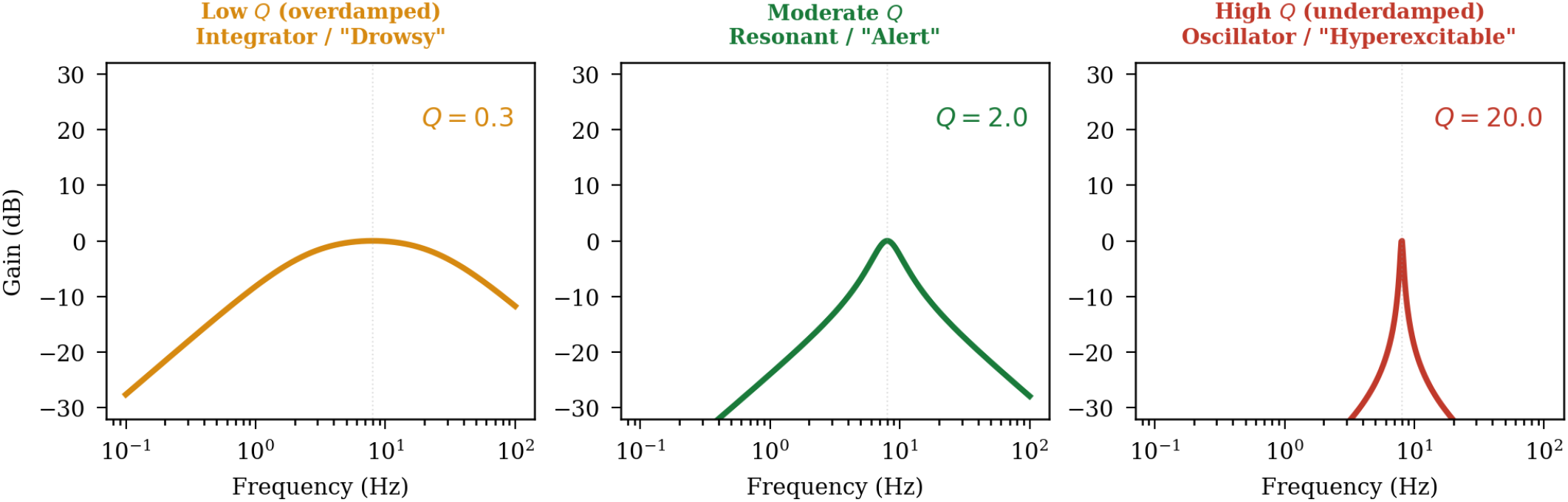
Neuromodulation as Q-factor control. Same RLC circuit at three Q values. Low Q: integrator mode (drowsy). Moderate Q: resonant, frequency-selective (alert). High Q: narrow oscillator (hyperexcitable).

**Fig. 5.**
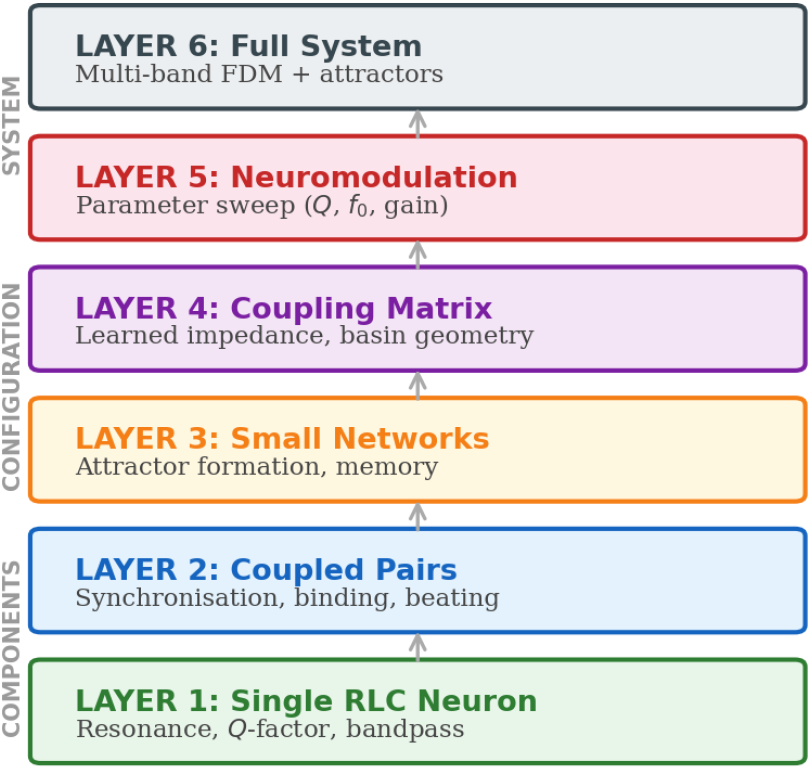
The full computational stack: from RLC components to analog computation. Each layer builds on the one below.

The answer is the attractor. The program is the impedance network.

## VIII. The Linearisation Gap

The RLC description is derived from linearisation of the Hodgkin–Huxley equations at subthreshold potentials. Impedance is a linear concept, valid for small perturbations around a fixed operating point [14]. This creates a gap when the framework extends to strongly nonlinear phenomena— spiking, attractor switching, chaos.

### A. Where the Framework Is Rigorous

Subthreshold potentials within ∼ 10–15 mV of rest, small-signal perturbations, timescales where ionic conductances remain approximately at their steady-state activation curves. In this regime, the neuron IS a parallel RLC circuit in the same sense that any system obeying a second-order linear ODE is an RLC circuit.

### B. Where It Becomes Heuristic

The effective RLC parameters are voltage-dependent: *L*_*eff*_ changes with gating variables, and at sufficiently hyperpolarised potentials the inductive term can vanish [21]. Spike generation is a highly nonlinear process violating the small-signal assumption. Network-level attractor dynamics involve bifurcations that cannot be derived from the linearised model.

### C. Bridging the Gap

Three levels of “resonance” must be distinguished because they dissociate at the linearisation boundary. *Subthreshold impedance resonance*—a peak in |*Z*(*f* )| —is the direct prediction of the RLC model and is rigorous within the small-signal regime. *Spike-output resonance*—frequency-selective firing probability and precision—requires the threshold nonlinearity and is most pronounced at low firing rates. *Information-transfer resonance*—band-pass filtering of stimulus information in the spike train—can remain low-pass in purely linear subthreshold dynamics and becomes band-pass only when spiking contributes. The framework’s Layers 1–2 concern subthreshold impedance resonance; the computational claims of Layers 3–6 depend on spike-output and information-transfer resonance, which require the bridge described next.

The subthreshold RLC dynamics set the *conditions* for computation: which frequencies are amplified, which inputs are selected, what phase relationships are established. The non-linear dynamics *execute* the computation: spikes fire, attractors are reached, patterns are completed. The connection is that nonlinear events are triggered by and timed relative to the linear dynamics. This connection is strongest when inputs are near-threshold and temporally structured—precisely the high-conductance state characterising in vivo cortical operation [5]. A neuron receiving strongly suprathreshold drive will spike at the input frequency regardless of its resonance; the RLC dynamics govern spike timing most powerfully in the near-threshold regime.

This is the standard relationship between small-signal and large-signal analysis in nonlinear circuit theory: small-signal parameters (gain, bandwidth, impedance) determine the operating regime and sensitivity; large-signal behaviour (saturation, switching, oscillation) determines the output. Both are needed; neither is sufficient alone.

## IX. Noise, Precision, and the Limits of Analog Computation

Analog computation is noise-limited. The dominant noise source in vivo is background synaptic bombardment: *σ*_*syn*_ ≈ 2–5 mV [5]. Using *σ*_*syn*_ ≈ 3 mV as representative:

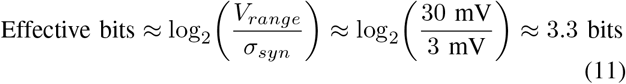

This is low by digital standards but is precisely the regime where Sarpeshkar [29] showed analog computation is optimal: at SNR below ∼ 20 dB, analog processing achieves higher energy efficiency than digital. The brain’s ∼ 3–4 effective bits per neuron, compensated by massive parallelism ( ∼ 10^10^ neurons), is the “noisy but efficient” regime.

### Phase precision is limited

For a theta oscillation ( ∼5 mV dendritic amplitude, ∼ 3 mV noise): Δ*ϕ*_*min*_ ≈ arcsin(*σ*_*syn*_*/A*) ≈ 37, giving ∼ 10 discriminable phase bins. At the soma in vivo, theta amplitude is often only ∼ 1–2 mV, making single-cell phase discrimination much worse. The claimed phase-coding capacity therefore requires population averaging, not single-cell readout. For a population of *n* neurons with independent noise, phase precision improves as 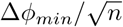 achieving ∼ 10° precision (36 discriminable bins) from somatic signals would require *n* ≈ (37*/*10)^2^ ≈ 14 neurons—biologically feasible within a single theta cycle given that hippocampal place cell ensembles typically involve tens to hundreds of co-active cells.

### Attractor basins must be noise-robust

The Hopfield capacity *P*_*max*_ ≈ 0.14*N* was derived for noiseless networks; at finite noise, basins shrink and effective capacity is lower [7].

## X. Correlative Versus Causal Evidence

### Causal

Subthreshold resonance (pharmacological block abolishes it; [15]). CPG reciprocal inhibition (silencing one half-centre stops rhythm; [24]). Theta frequency constraining WM capacity (tACS at different frequencies shifts capacity; [27], [28]).

### Correlative

Phase-binding/synchrony (co-occurrence during binding tasks, but most studies measure correlation not causation; [6]). Cross-frequency coupling and information content (PAC predicts performance but does not establish low frequencies causally driving high-frequency content; [3]). Gamma clocking of items (gamma cycle durations vary, undermining strict periodic clocking; [17]).

The framework’s claims are therefore graded: strongly supported (resonance, CPG inhibition), moderately supported (attractor dynamics, neuromodulatory parameter tuning), and provisionally supported (phase binding, gamma clocking). The phase-based Hebbian rule (Equation 10), the RLC advantage for attractor dynamics, and the neuromodulatory Q-sweep predictions are theoretical and await empirical testing.

## XI. Falsifiability

### Resonance is computationally relevant

Confirmed in entorhinal stellate cells [9]; independent tests in inferior olive and thalamic relay neurons would extend this. Quantitatively: neurons with higher *Q* should show lower spike-timing jitter at *f*_0_, with the predicted relationship *σ*_*timing*_ ∝ 1*/Q*.

### Phase encodes relational information

Falsified if disrupting phase-locking (e.g., via independent tACS to two populations) does not impair binding. Specifically: the binding-failure threshold should scale with coupling strength as Δ*f*_*crit*_ = *g/*(2*πC*_*m*_) (from Equation 6).

### Attractor dynamics implement retrieval

Falsified if trajectories from similar initial conditions diverge rather than converge in a recurrent network with Hebbian connectivity. The RLC framework additionally predicts that retrieval speed scales with *Q*: higher-*Q* networks should show faster convergence (fewer oscillatory transients before settling) in deep attractor basins.

### Neuromodulation operates as parameter tuning

Falsified if serotonin shifts *f*_0_ upward (predicted: downward) or ACh broadens tuning (predicted: sharpens). Note that this prediction applies to cells where *I*_*h*_ is the dominant resonance current; in thalamocortical neurons where *I*_*T*_ dominates [35], the neuromodulatory mapping may differ.

### RLC outperforms RC

Build both in neuromorphic hardware; if the RC circuit matches RLC performance on tasks requiring frequency selectivity, phase coding, or temporal precision, the framework’s added complexity is unjustified.

## XII. Discussion

### A. Relationship to Existing Frameworks

The framework builds on established foundations: Koch [16] on biophysical cable theory, Sarpeshkar [29] on the energy optimality of analog neural computation, Hopfield [11] on attractor dynamics, Lisman and Jensen [17] on theta-gamma coding, and Aoyagi [1] and Scarpetta et al. [30] on oscillatory attractor networks with phase-coded learning. The present contribution is to connect these into a single stack grounded in measurable impedance quantities—*f*_0_, *Q*, and *L*_*eff*_ —that constrain which computations are physically achievable in a given neural population.

### B. What the Framework Adds

Three contributions distinguish this work. First, the *phase-based Hebbian rule* (Equation 10) grounds synaptic learning in impedance structure: the phase relationships being learned are set by the resonant properties of the neurons, not free parameters. Second, the *attractor taxonomy* connects fixed-point, limit-cycle, and chaotic attractors to RLC dynamics and neuromodulatory Q-factor control, constraining oscillator parameters to biophysical measurements rather than treating them as free [1]. Third, the explicit *regime of validity* analysis (Sections VIII–X) makes the framework’s assumptions transparent and its claims testable.

### C. Implications for Neuromorphic Engineering

The framework provides a principled design library for neuromorphic systems. Rather than approximating neural dynamics with digital spiking neural networks, neuromorphic engineers can implement the same analog circuits—in the same topology, operating in the same signal-processing regime. The key insight is that the computation resides in the continuous dynamics, not in the spikes. A neuromorphic chip preserving these dynamics (RLC silicon neurons, analog coupling, continuous-time operation) will capture computational properties that digital approximations cannot [19], [23]. Subthreshold CMOS implementations exploit exponential transistor characteristics for low power, but scaling large networks requires managing device mismatch and temperature sensitivity. The framework predicts computation in the low-SNR regime ( ∼ 3.3 effective bits) where analog processing has an energy advantage over digital [29] and where device mismatch is partially absorbed by the noise tolerance the biological system already requires. Chip-in-the-loop training— learning algorithms operating on physical hardware, adapting around device-specific offsets—is likely necessary for any implementation of Layers 3–6.

## XIII. Conclusion

We have presented a six-layer computational framework building from single-neuron RLC resonance to system-level attractor dynamics. The framework is grounded in measurable electrical quantities, generates testable predictions, and explicitly states its regime of validity and the evidential status of each claim.

The brain computes by being an analog circuit. The tools to understand that computation—impedance spectroscopy, transfer functions, stability analysis, noise theory—have existed in electrical engineering for a century. This paper argues that applying them systematically to neural circuits, with the rigour and humility that both the circuits and the tools deserve, opens a complementary perspective on neural computation that captures dynamics rate-based models miss.

## Notes

### Competing Interest Statement

The authors have declared no competing interest.

